# Spatiotemporal Distribution of Vector Mosquito Species and Areas at Risk for Arbovirus Transmission in Maricopa County, Arizona

**DOI:** 10.1101/2022.07.28.501907

**Authors:** André B. B. Wilke, Dan Damian, Maria Litvinova, Thomas Byrne, Agnese Zardini, Piero Poletti, Stefano Merler, John-Paul Mutebi, John Townsend, Marco Ajelli

## Abstract

Mosquito-borne diseases are a major global public health concern and mosquito surveillance systems are essential for the implementation of effective mosquito control strategies. The objective of our study is to determine the spatiotemporal distribution of vector mosquito species in Maricopa County, AZ from 2011 to 2021, and to identify the hotspot areas for West Nile virus (WNV) and St. Louis Encephalitis virus (SLEV) transmission in 2021. The Maricopa County Mosquito Control surveillance system utilizes BG-Sentinel and EVS-CDC traps throughout the entire urban and suburban areas of the county. We estimated specific mosquito species relative abundance per unit area using the Kernel density estimator in ArcGIS 10.2. We calculated the distance between all traps in the surveillance system and created a 4 km buffer radius around each trap to calculate the extent to which each trap deviated from the mean number of *Cx. quinquefasciatus* and *Cx. tarsalis* collected in 2021. Our results show that vector mosquito species are widely distributed and abundant in the urban areas of Maricopa County. A total of 691,170 *Culex quinquefasciatus*, 542,733 *Culex tarsalis*, and 292,305 *Aedes aegypti* were collected from 2011 to 2022. The relative abundance of *Ae. aegypti* was highly seasonal peaking in the third and fourth quarters of the year. *Culex quinquefasciatus*, on the other hand, was abundant throughout the year with several regions consistently yielding high numbers of mosquitoes. *Culex tarsalis* was abundant but it only reached high numbers in well-defined areas bordering natural and rural areas. We also detected high levels of heterogeneity in the risk of WNV and SLEV transmission to humans disregarding traps geographical proximity. The well-defined species-specific spatiotemporal and geographical patterns found in this study can be used to inform vector control operations.

## Introduction

Mosquito-borne diseases are a major global public health concern [1–4]. The incidence of dengue has substantially increased globally in the last decades, from 500,000 reported cases in 2000 to 2.4 million in 2010 and 5.2 million in 2019 [5]. The current estimate is 390 million dengue virus (DENV) infections worldwide every year [6]. Between 2015 and 2018, there were more than 1 million confirmed cases of Zika in the Americas followed by a significant increase in fetus malformation as a direct consequence of Zika virus (ZIKV) infection during pregnancy [3,4].

Albeit on a smaller scale, arboviral infections have also increased in the United States. The ZIKV was introduced multiple times in the United States in 2016 leading to 224 locally transmitted human cases in Florida and Texas [9–11]. Local transmission of the DENV has also been reported in Florida. In 2020, Monroe County experienced a major dengue outbreak with 65 confirmed locally transmitted cases [12–14]. In 2022, 33 locally transmitted cases of dengue were reported in Miami-Dade County [12].

While local outbreaks of dengue and Zika were reported in the United States, the West Nile virus (WNV) has become endemic and the most widespread mosquito-borne arbovirus in North America [15]. WNV has been detected in more than 60 mosquito species in the continental United States (CONUS) [16]; however, selected species in the genus *Culex* such as *Culex quinquefasciatus, Culex pipiens, Culex tarsalis*, and *Culex nigripalpus* are responsible for driving most of the epizootic and epidemic WNV transmission. Birds are the natural reservoir hosts of WNV as they are able to develop viremia high enough to infect mosquito vectors. Humans, horses, and most mammals are not able to amplify the virus to viremic levels high enough to infect mosquitoes and are therefore considered dead-end hosts [16–18].

Since it was first detected in New York in 1999 [19], the WNV has progressively become endemic in all states in the contiguous United States and has been sporadically detected in Alaska, Hawaii, and Puerto Rico [20]. From 1999 to 2020 a total of 52,532 human cases were reported in the United States leading to 2,456 deaths [21].

Maricopa County, Arizona, is located in the Sonoran Desert ecoregion spreading through 23,890 km^2^. It has a hot desert climate with long hot summers and short winters [22]. According to the 2020 census [23], Maricopa County has a population of 4,420,568 people, being the most densely populated county in Arizona as well as in the Sonoran Desert ecoregion. There are three major cities in Maricopa County, Phoenix (the Arizona state capital, and the fifth-most populous city in the United States), Mesa, and Chandler. It has historically been one of the counties most affected by West Nile virus disease in the United States with an average of 70 neuroinvasive cases reported every year in the last decade (2010 to 2020). However, 1,427 neuroinvasive cases of West Nile were reported in this county in 2021, the highest ever reported in a single county in the United States [12].

Controlling mosquito populations is widely considered the most effective way of preventing the spread of mosquito-borne diseases [24]. However, the development and implementation of effective mosquito control strategies in urban areas are complex and difficult to put into action [25]. They rely on a framework consisting of multiple successive actions that need to be implemented for a successful intervention. The first step in the implementation of effective strategies is the development of an extensive and reliable mosquito surveillance system [26]. Such a system is key to assessing the mosquito community composition, identifying factors influencing the population dynamics of mosquito species, estimating the relative abundance of populations of different vector mosquito species, and providing evidence for ascertaining and anticipating patterns in mosquito population dynamics and demographics that are essential for the development of mosquito control strategies. The aim of this study is twofold: (i) determine the spatiotemporal distribution of mosquito species vectoring different arboviruses (including WNV, St. Louis Encephalitis virus – SLEV, DENV, ZIKV) in Maricopa County, Arizona, from 2011 to 2021; and (ii) investigate the hotspot areas for WNV and SLEV transmission in 2021.

## Methods

The Maricopa County Environmental Services Department Vector Control Division surveillance grid covers the entire urban and peri-urban areas of the county with approximately one trap per square mile. The surveillance system is comprised of BG-Sentinel traps baited with BG-Lure (Biogents AG, Regensburg, Germany) (ranging from 13 traps in 2016 to 8 traps in 2021) and EVS-CDC traps baited with CO_2_ (ranging from 478 traps in 2011 to 825 traps in 2021). Each trap is deployed once a week for 24 hours for 50 weeks a year (Figure 1). All collected mosquitoes are transported to the Maricopa County Environmental Services Department Vector Control Division Laboratory and morphologically identified to species using taxonomic keys [27]. BG-Sentinel traps baited with BG-Lure and EVS-CDC traps baited with CO_2_ attract female mosquitoes seeking hosts for blood-feeding [28]. Even though male mosquitoes were sometimes collected, male data was not included in the proposed analyses.

**Figure 1.**
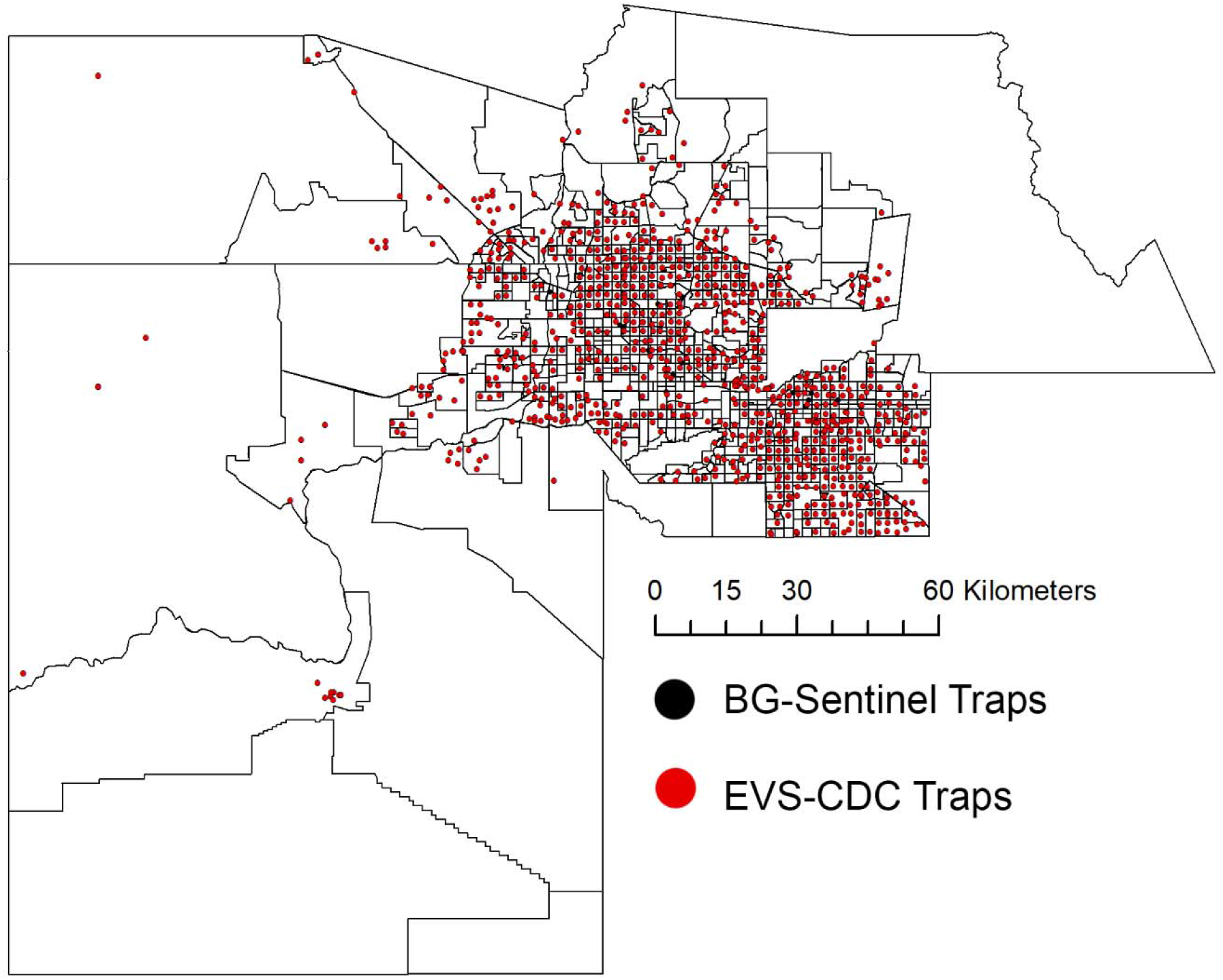
Map of Maricopa Mosquito Control Surveillance System. BG-Sentinel traps are shown in black; EVS-CDC traps are shown in red.

For 2021, virological surveillance was done for all collected *Cx. quinquefasciatus* and *Cx. tarsalis* mosquitoes in pools of up to 50 female mosquitoes. RNA was extracted using MagMAX extraction kit (Thermo Fisher Scientific). Samples were prepared in a PCR hood (Air Clean 600 PCR workstation “Reagent Prep Hood”). Each PCR reaction included WNV and SLEV positive and negative controls (Table 1).

**Table 1.**
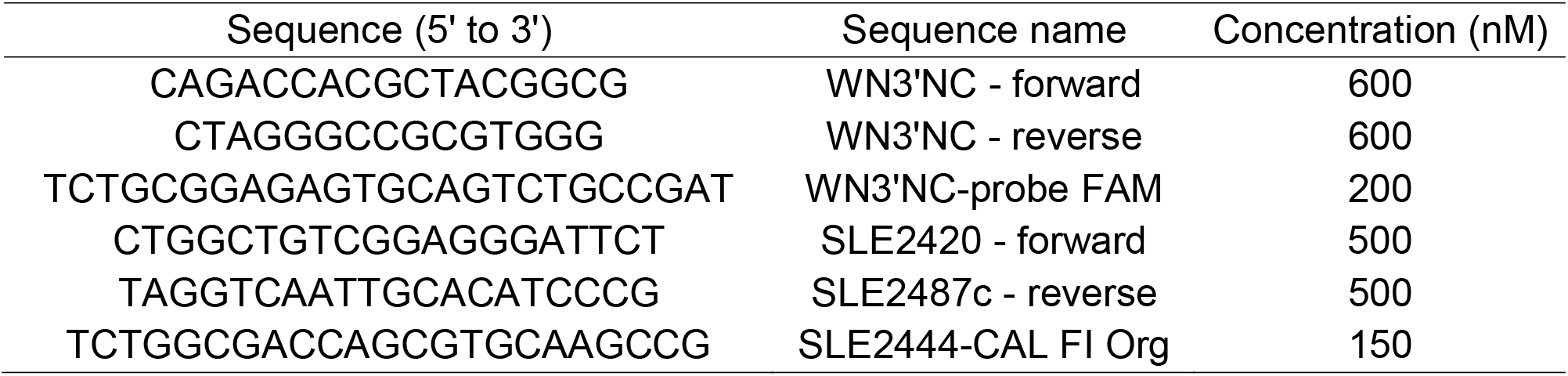
West Nile and St. Louis Virus Real-Time RT-qPCR primers.

Reactions were carried out using Taqman Master Mix (Thermo Fisher Scientific) on a QuantStudio 5 at 50°C for 5 minutes, 95°C for 20 seconds, and 40 cycles of: 95°C for 3 seconds and 60°C for 30 seconds. Samples were considered positive when Ct values were below 34 and negative when qPCR reaction resulted in no detectable Ct value after 40 cycles.

We used the geodesic method for densities in square kilometers to calculate the magnitude-per-unit area (number of inputs per point - i.e., mosquito relative abundance) using the Kernel density estimator in ArcGIS 10.2 (Esri, Redlands, CA). Figures were created with layers available at the Maricopa County GIS Mapping Applications available at: https://www.maricopa.gov/3942/GIS-Mapping-Applications.

We calculated the distance between all traps in the surveillance system (linear distance in meters) and created a 4 km buffer radius around each target trap. Traps within each buffer with no data were removed from the analysis. We also removed outliers in each buffer by excluding observations that lie outside the expected range of the variability of the mean 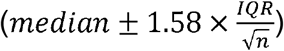 to calculate mean buffer values. If the target trap was among the removed outliers, it was identified as a hot (cold) spot if the number of collected mosquitoes was above (below) the buffer mean (Figure 5). Hot/cold spot traps were defined to highlight unexpectedly high/low number of collected mosquitoes relative to the traps in the same area - information useful for the design of tailored mosquito control interventions. This analysis was conducted in R (version 4.2.2).

To assess the risk of arbovirus transmission posed by the abundance of vector mosquito species, we used the 2021 data on mosquito vector species presence, density, and infection rate to calculate the Vector Index (VI: average number of mosquitoes collected by trap night multiplied by the infection rate) [29]. The infection rate was estimated as the ratio of the mosquito pools that tested positive for WNV or SLEV over the total number of tested pools. Then, we used ArcGIS to create a 1-km buffer around each individual trap and extracted the 2020 total population from the U.S. Census using the weighted centroid geographic retrieval methodology [30].

## Results

### Spatiotemporal Distribution of Vector Mosquito Species in Maricopa County, Arizona

The Maricopa County Environmental Services Department Vector Control Division Surveillance System was comprised of 478 traps in 2011 and new traps were gradually included over the years as the surveillance system expanded to 833 traps in 2021. From 2016 to 2021 the number of traps remained nearly constant. A total of 691,170 *Cx. quinquefasciatus*, 542,733 *Cx. tarsalis*, and 292,305 *Aedes aegypti* were collected from 2011 to 2022 by the traps in the Surveillance System (Figures 2 and 3, Supplementary Table 1).

**Figure 2.**
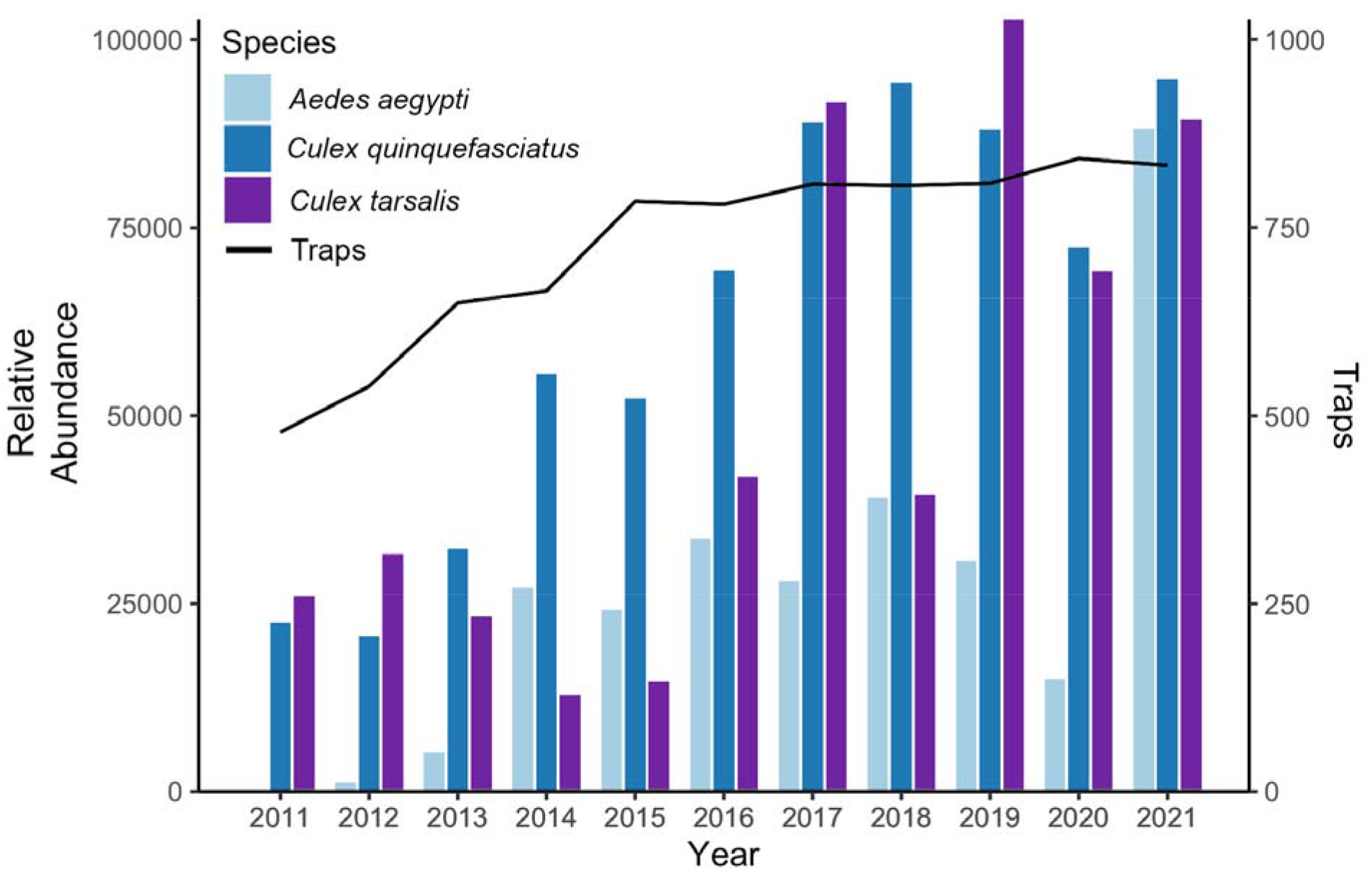
Number of active traps in the Maricopa County Environmental Services Department Vector Control Division Surveillance System (black line) and the number of mosquitoes collected by year from 2011 to 2021.

**Figure 3.**
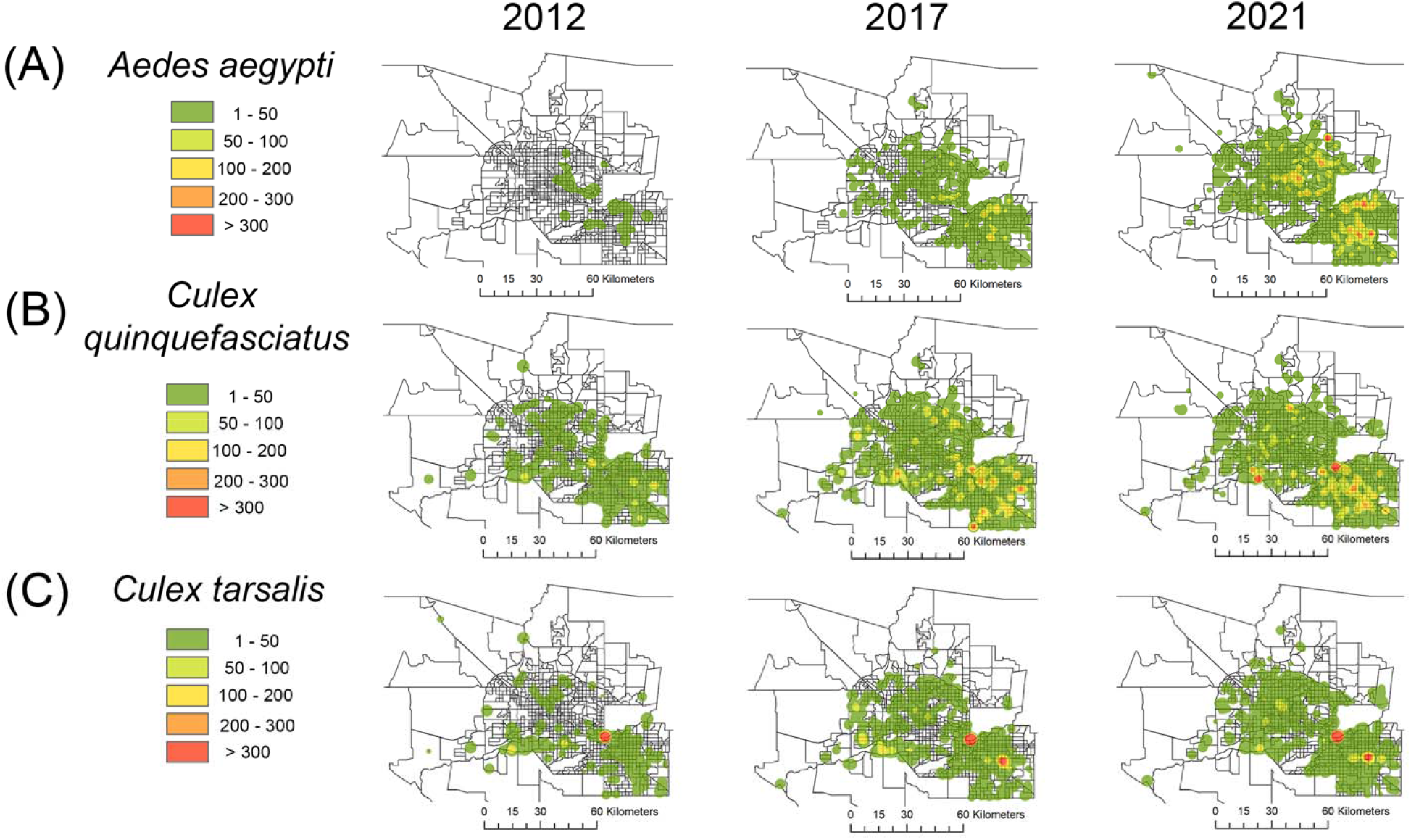
Kernel density-based heat map of the relative abundance of vector mosquito species collected by the Maricopa County Environmental Services Department Vector Control Division Surveillance System in 2012, 2017, and 2021. The color gradient represents the sum of mosquitoes collected by each trap in a 12-month period; (A) *Aedes aegypti*; (B) *Culex quinquefasciatus*; and (C) *Culex tarsalis*. The figure was produced using ArcGIS 10.2 (Esri, Redlands, CA), using layers available at the Maricopa County GIS Mapping Applications available at: https://www.maricopa.gov/3942/GIS-Mapping-Applications.

Over the study period, *Ae. aegypti* gradually expanded its range and relative abundance in the urban areas of Maricopa County. In particular, compared to 2020, a substantial increase in the range of *Ae. aegypti* was observed in 2021 when a total of 88,187 specimens were collected by the traps in the surveillance system (Figure 3A and Supplementary Figure 1A). In contrast to *Ae. aegypti, Cx. quinquefasciatus* was already commonly found in the urban areas in Maricopa County in 2011. However, since 2011, *Cx. quinquefasciatus* became more confined to specific areas of the county and was particularly abundant in those areas. Since 2016, *Cx. quinquefasciatus* started to increase in abundance, especially in the southeastern region of Maricopa County (Figure 3B and Supplementary Figure 1B). Although *Cx. tarsalis* could be detected in urban areas, this species was mainly confined to specific areas of Maricopa County, primarily to areas inside or bordering natural or rural areas. By 2021, *Cx. tarsalis* had gradually expanded its range, but it remained highly abundant in this ecological niche (Figure 3C, Supplementary Figure 1, Supplementary Tables 2 and 3).

Our results show great seasonal variations in the mean numbers of mosquitoes collected in Maricopa County. Rainfall is one of the possible drivers for the proliferation of mosquitoes in the region, and the peaks in abundance of *Cx. quinquefasciatus* and *Cx. tarsalis* overlap with the peak of the rainy season in the Spring and later in early Fall. Furthermore, rainfall is potentially an important driver for the increase in the abundance of *Ae. aegypti* over the year until the beginning of the dry season, when the abundance of *Ae. aegypti* populations is drastically reduced. Each of the three vector mosquito species, *Ae. aegypti, Cx. tarsalis*, and *Cx. quinquefasciatus*, displayed different population dynamic patterns peaking in abundance at different times of the year. This indicates that even though rainfall is an important driver for their proliferation, each species responded differently to the rainfall (Figure 4).

**Figure 4.**
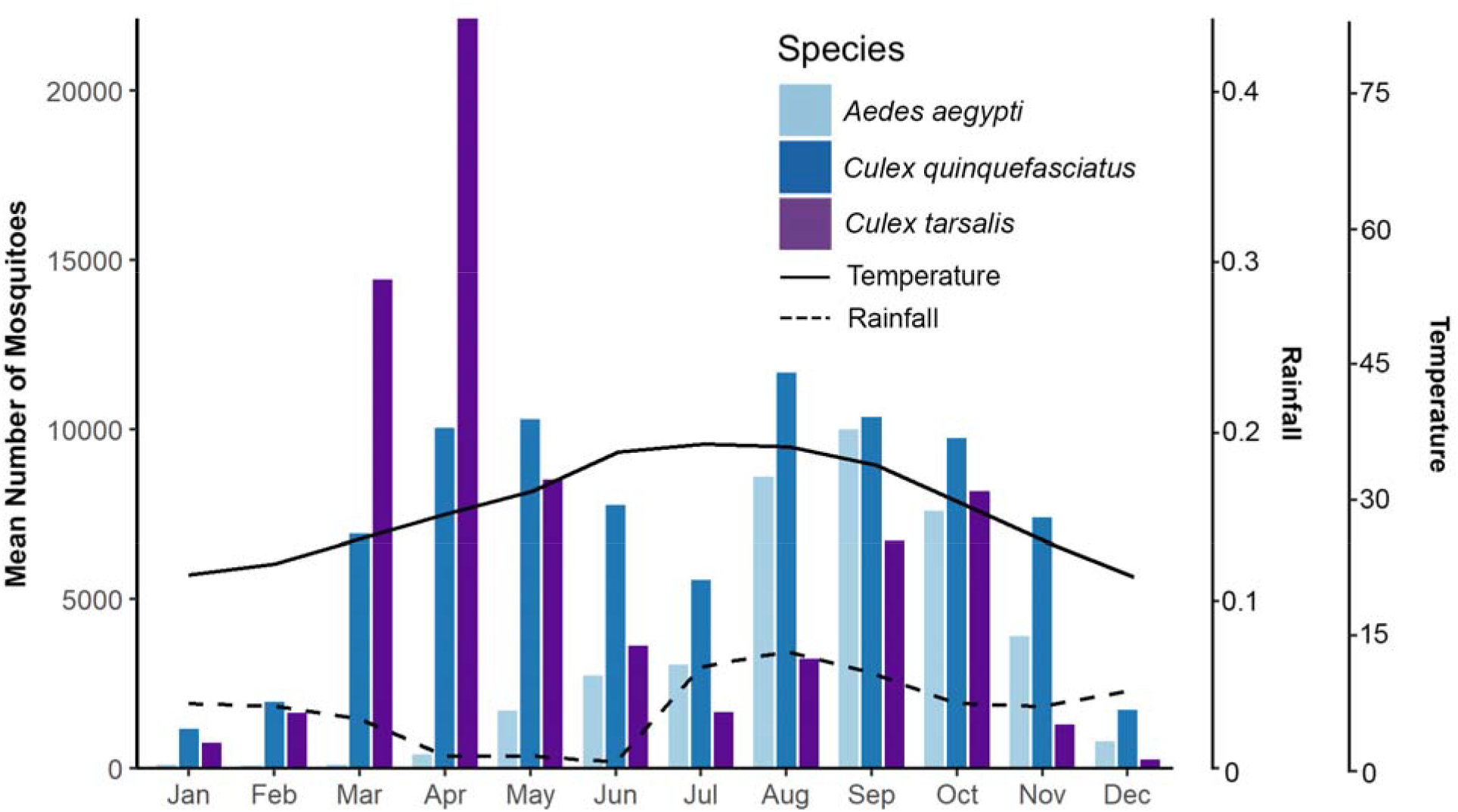
Mean number of *Aedes aegypti, Culex quinquefasciatus*, and *Culex tarsalis* collected by the Maricopa County Environmental Services Department Vector Control Division Surveillance System from 2016 to 2021. Solid line represents the monthly mean average temperature (Celsius) between 2016 and 2021; Dashed line represents the monthly mean average rainfall (centimeters) between 2016 and 2021.

The mean number of collected *Ae. aegypti* varied greatly within each year. It was virtually absent during the first quarter of the year yielding a low number of mosquitoes collected by the traps in the surveillance system. It then gradually increased in abundance and range during the second quarter, showing the highest abundance in the southeastern part of the county. This species peaked in abundance during the third quarter of the year covering most of the urban areas of Maricopa County and reaching high abundance in several geographical areas. The mean number of *Ae. aegypti* collected in the fourth quarter of the year was smaller than in the third quarter, albeit areas with high abundance were still present (Figure 5A).

**Figure 5.**
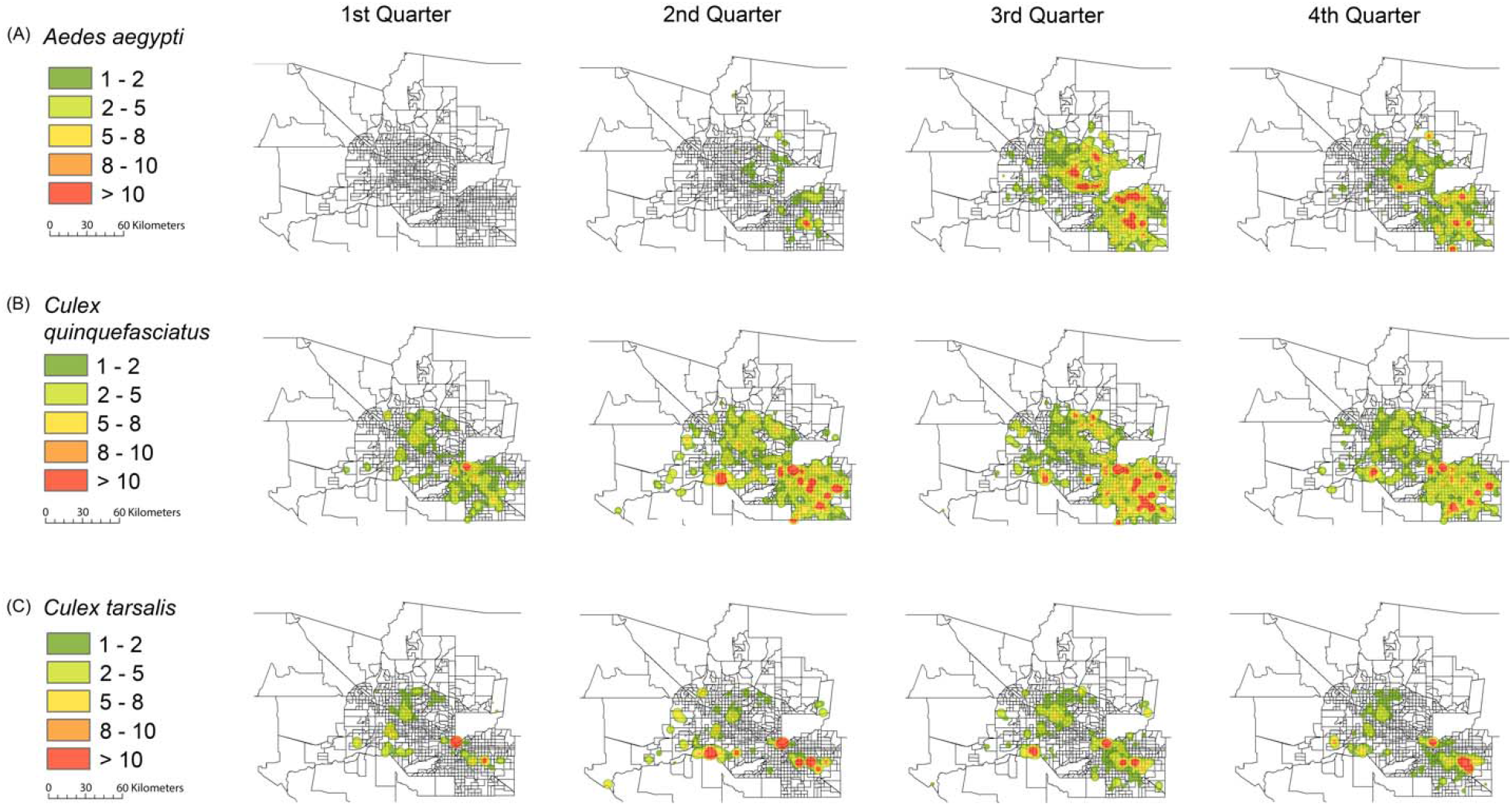
Kernel density-based heat map of the mean number of *Aedes aegypti, Culex quinquefasciatus*, and *Culex tarsalis* collected by the Maricopa County Environmental Services Department Vector Control Division Surveillance System from 2016 to 2021. The color gradient represents the mean number of mosquitoes collected by each trap in a 3-month period. The figure was produced using ArcGIS 10.2 (Esri, Redlands, CA), using layers available at the Maricopa County GIS Mapping Applications available at: https://www.maricopa.gov/3942/GIS-Mapping-Applications.

*Culex quinquefasciatus* was abundantly found in Maricopa County year-round. The mean number of *Cx. quinquefasciatus* was the highest during the second and third quarters of the year with many areas of the county yielding an average higher than 10 mosquitoes per trap per day. One region bordering rural and leisure areas in the southeastern part of the county yielded more than 10 mosquitoes per trap per day on average year-round (Figure 5B). *Culex tarsalis* was moderately affected by seasonality. The mean number of *Cx. tarsalis* remained constant during the second, third, and fourth quarters of the year with specific areas yielding high numbers of mosquitoes. During the first quarter, collections of *Cx. tarsalis* were not as abundant as in the other quarters of the year, but for one region bordering rural and leisure areas in the southeastern part of the county. Furthermore, *Cx. tarsalis* yielded the highest average number of mosquitoes collected in a selected group of traps bordering natural and rural areas. Even though the mean number of *Cx. tarsalis* reached high levels in those specific areas, it was not abundantly found in urban areas. (Figure 5C).

### Hotspot Areas for the transmission of WNV and SLEV in Maricopa County, Arizona

To further our understanding and provide operational input in case of a WNV and SLEV arbovirus outbreak in Maricopa County, we focused on the mosquito and arbovirus surveillance data from 2021, when Maricopa County reported a record-breaking number of neuroinvasive WNV cases. To identify highly conducive environments for the proliferation of *Cx. quinquefasciatus* and *Cx. tarsalis*, we created a 4 km buffer radius around each trap (corresponding to 15 traps on average, range: 1-34). Then, we calculated the number of traps in each of the buffers, and the total and mean number of *Cx. quinquefasciatus* and *Cx. tarsalis* collected by the traps in the cluster. Our results show that from the 820 traps that collected *Cx. quinquefasciatus* female mosquitoes in 2021, 90 were above their respective buffer average, with 21 influential points (defined as traps yielding an average number of mosquitoes at the 97.5^th^ percentile of each of the clusters). From the 754 traps that collected *Cx. tarsalis* female mosquitoes in 2021, 76 were above their respective buffer average, with 19 influential points (Figure 6).

**Figure 6.**
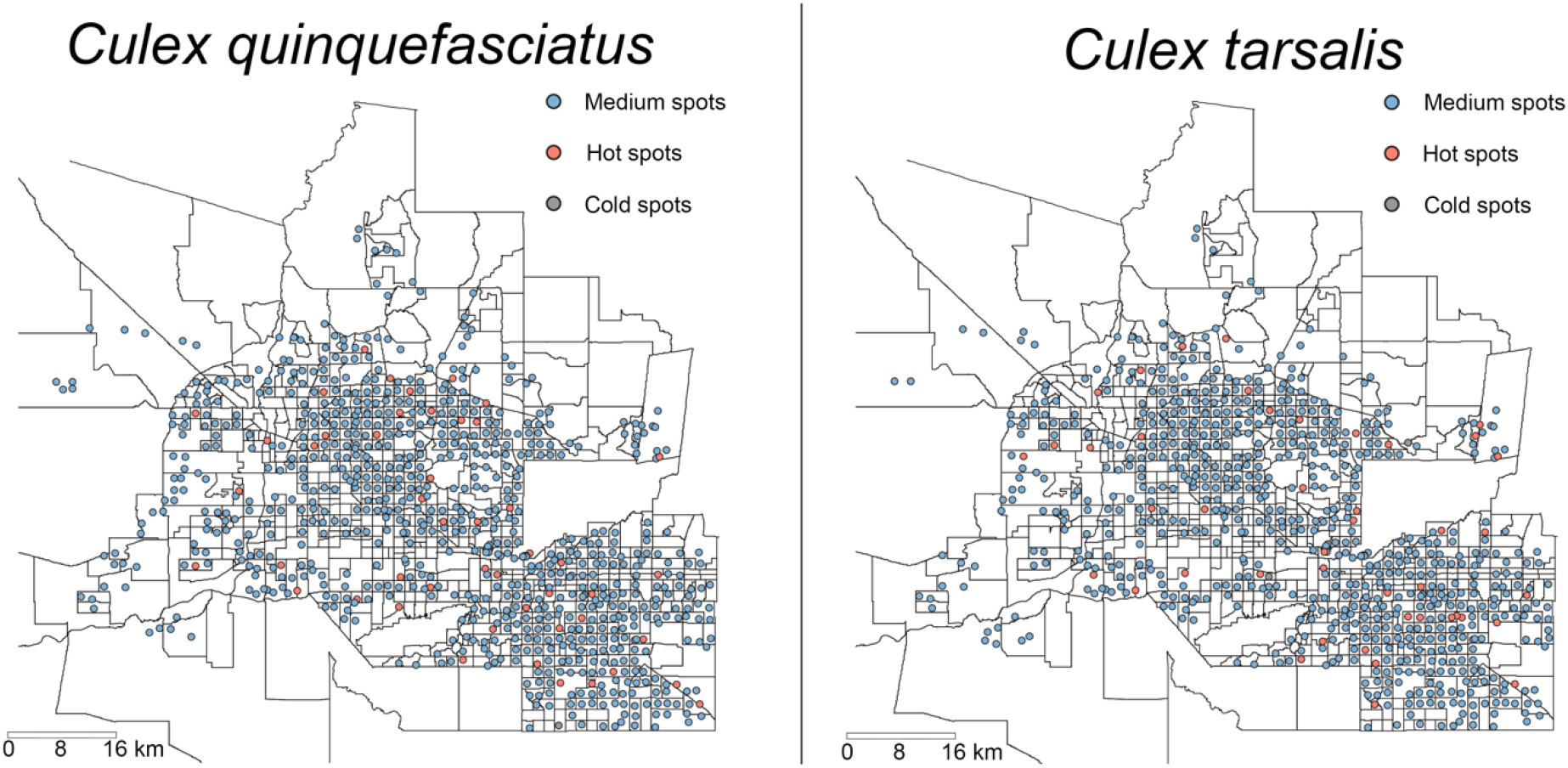
Hot spot and cold spot traps that collected a higher (red) or lower (gray) number of female mosquitoes *Cx. quinquefasciatus* and *Cx. tarsalis* relative to the traps in their respective 4 km buffer radius in Maricopa County, Arizona in 2021.

To assess the locations in Maricopa County, Arizona, with the highest WNV and SLEV disease transmission risk, we calculated the VI. We then created a map and overlaid the VI with human population density. Identifying locations with high human population densities and high levels of infected mosquitoes serves as a powerful indicator of the highest-risk locations for WNV and SLEV transmission.

Our results show high levels of heterogeneity in the risk of WNV and SLEV transmission to humans disregarding the geographical proximity of the traps (i.e., the risk of arbovirus transmission has been driven locally at the microgeographic scale and varied greatly between closely located traps). In 2021, WNV was detected in *Cx. quinquefasciatus* mosquito pools at least once in 304 different traps, with VI values ranging from 0.02 to 0.12, and in *Cx. tarsalis* mosquito pools at least once in 222 different traps, with VI values ranging from 0.02 to 0.1. SLEV was detected in *Cx. quinquefasciatus* mosquito pools at least once in 15 different traps. It was also detected at least once in 24 different traps in *Cx. tarsalis* mosquito pools. In both cases, the VI values ranged from 0.02 to 0.04 (Figure 7).

**Figure 7.**
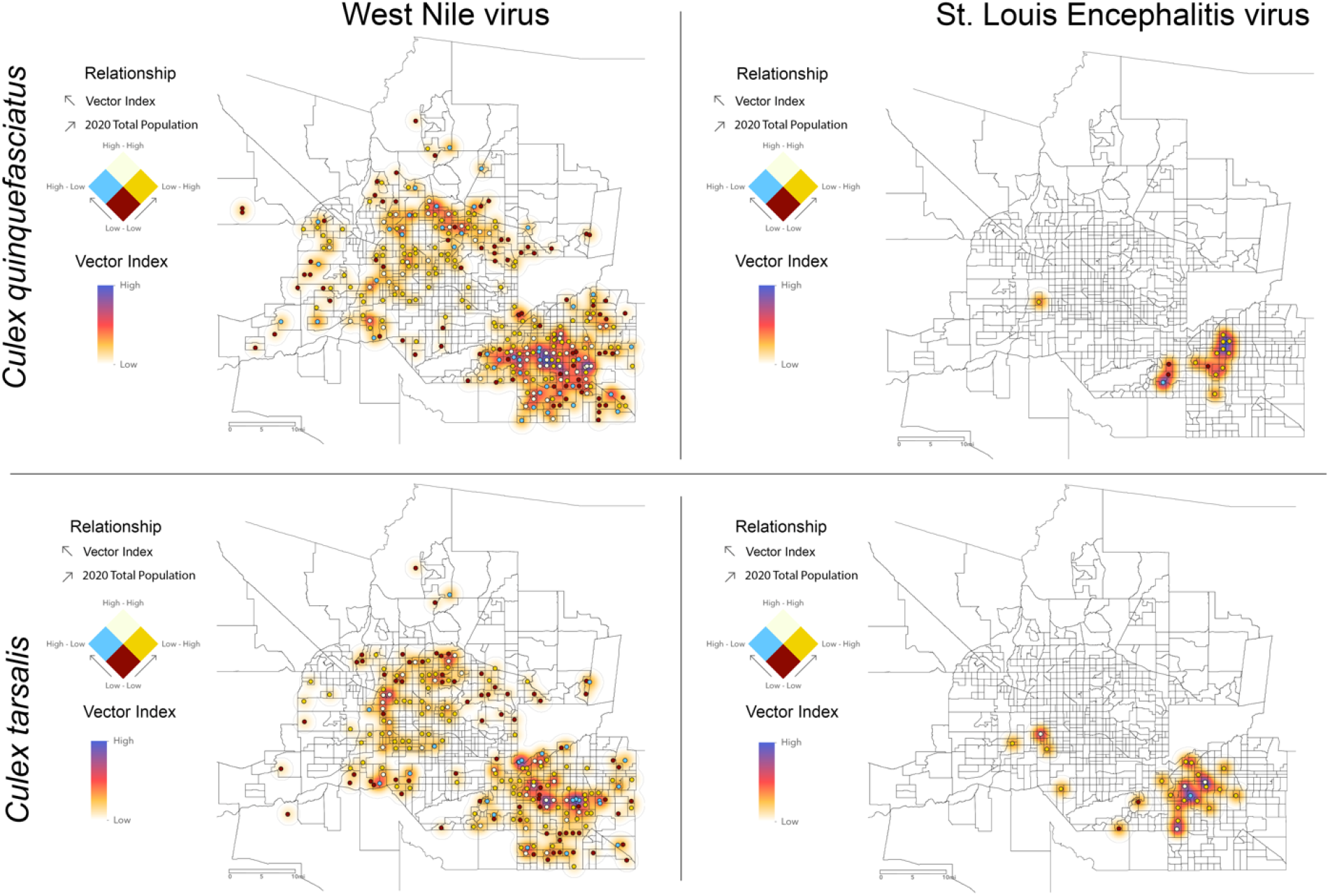
High-risk areas for the transmission of WNV and SLEV in Maricopa County, Arizona in 2021. Relationship between the estimated average number of WNV and SLEV infected *Cx. quinquefasciatus* and *Cx. tarsalis* by trap night (Vector Index) in Maricopa County, Arizona, in 2021, and the 2020 total population from the U.S. Census in a 1-km buffer around each individual trap. White dots represent areas with high Vector Index and high human population density, blue dots represent areas with high Vector Index and low human population density, yellow dots represent areas with low Vector Index and high human population density, and dark red dots represent areas with low Vector Index and low human population. The overall heterogeneity of the Vector Index for WNV and SLEV circulating in *Cx. quinquefasciatus* and *Cx. Tarsalis* populations are represented using a heatmap.

## Discussion

Anthropogenic changes in the environment create favorable conditions for the development of vector mosquito species that can exploit the resources available in urban areas [31]. The decreased biodiversity in those urban areas increases the contact rate between mosquito vector species and human hosts, increasing the risk of arbovirus transmission to humans [14,32,33]. Our results show that vector mosquito species are widely distributed and abundantly found in the urban areas of Maricopa County, Arizona.

*Aedes aegypti* gradually become more abundant and expanded its range since only one specimen was collected in Maricopa County in 2011. In 2012, 1,189 *Ae. aegypti* were collected; by 2014 that number increased to 27,208, and by 2021 a total of 88,187 specimens were collected indicating a substantial increase in its relative abundance over the years. A similar phenomenon could be observed for *Cx. tarsalis* and *Cx. quinquefasciatus*, and although they were already present in great numbers since 2011, with more than 20,000 specimens from each species collected that year, reaching 89,401 collected specimens *of Cx. tarsalis* and 94,774 of *Cx. quinquefasciatus* in 2021. We should stress that mosquitoes were collected using BG-Sentinel and CDC traps, and new traps were gradually added during the period of this study increasing the sampling effort over time. Even though our results show an increase in the relative abundance of mosquitoes, the different sampling effort over the years remains a study limitation.

The high abundance of vector mosquitoes in Maricopa County represents a potentially dangerous scenario. *Aedes aegypti* is the primary vector of dengue, chikungunya, yellow fever, and Zika viruses, and *Cx. quinquefasciatus* and *Cx. tarsalis* are vectors of the WNV, Saint Louis Encephalitis, Eastern Equine encephalitis, and Japanese Encephalitis viruses [34]. Moreover, differently from dengue and other *Aedes*-borne diseases where humans do not represent dead-end hosts, for WNV and SLEV the risk of human cases directly depends on the number of individuals exposed to bites from infected mosquito vectors. Thus, the high concentration of human population in urban areas of Maricopa County living in close proximity to infected mosquito vectors has the potential to lead to major disease burden, as it was the case in 2021, when 1,427 neuroinvasive WNV cases were reported in the county [12].

The relative abundance of *Ae. aegypti* was highly seasonal with fewer mosquitoes collected in the first 3 months of the year followed by a moderate increase in numbers in the second quarter indicating that the environmental conditions were not conducive to the proliferation of *Ae. aegypti* during the first quarter of the year. However, as the year progresses, a substantial increase in the abundance of *Ae. aegypti* was observed in the third and fourth quarters. The observed seasonal variation in the abundance of *Ae. aegypti* can potentially be an important driver for arbovirus transmission in Maricopa County as its abundance and range increase as the year progresses, thus, potentially increasing the contact rate between *Ae. aegypti* and human hosts. *Culex quinquefasciatus*, on the other hand, was abundantly found during all seasons of the year. It was not as abundant in the first quarter as it was during the remaining of the year, but it could be found throughout the urban areas during that time of the year. Several regions consistently yielded high numbers of *Cx. quinquefasciatus* mosquitoes indicating that those areas were hot spots and were able to support high numbers of *Cx. quinquefasciatus* year-round. *Culex tarsalis* was also abundantly found in Maricopa County but in contrast to *Ae. aegypti* and *Cx. quinquefasciatus*, it only reached high numbers in well-defined areas bordering natural and rural areas. However, this is not surprising because *Cx. tarsalis* larval habitats include irrigation agricultural runoff, wetlands, sewage effluent, oil field run-off, and lake beds [35], and all these habitat types are rarely associated with urban areas. Human cases of WNV disease have been consistently reported in Arizona, being one of the most affected states in the United States. The high abundance of *Culex* vector species, especially the year-round presence of *Cx. quinquefasciatus*, has major implications for the likelihood of WNV transmission in Maricopa County [12]. In sum, our results showed well-defined spatiotemporal patterns for each of the species included in this study. However, further analyses are needed to investigate the main drivers for the variations in the population dynamics of each mosquito vector species that are essential for the development and implementation of effective source reduction and other mosquito control strategies [36].

The cluster analysis results aimed to detect influential point traps that collected a significantly higher number of either *Cx. quinquefasciatus* or *Cx. tarsalis* as compared to other traps in their proximity indicate that resources available at the microgeographic scale were responsible for their proliferation. Further analysis is needed to understand which local geographic features (e.g., urban parks, fishing ponds, sports complexes, as appears to be the case in our data) may have contributed to the observed patterns. Identifying such factors is instrumental to guide vector control operations.

The identification of high-risk areas for the transmission of WNV and SLEV is vital for the development of vector control strategies aiming at reducing the abundance of *Cx. tarsalis* and *Cx. quinquefasciatus* and thus the risk of infection. Our results show the overlap between areas with high human population densities and high levels of infected mosquitoes. These areas are priority targets for the development and implementation of vector mosquito control strategies. Furthermore, there is a need to assess and validate the effectiveness of different control methods and strategies aimed to reduce mosquito populations in urban areas. These activities could focus on how the proliferation of vector mosquito species can be locally reduced by source reduction, community engagement, and by modifying specific features of the urban built environment [36–39]. Developing and implementing effective mosquito control strategies is a difficult task with many factors that can potentially hinder the effectiveness of mosquito control interventions, such as levels of insecticide resistance, diel activity of target mosquito species, and outdated and ineffective mosquito control tools [40–42].

Our analysis of high-risk areas for WNV and SLEV infection considers only the density of the human population, disregarding other characteristics such as the population’s, age structure and underlying health status, as well as other social determinants including individuals’ socioeconomic status, and characteristics of the neighborhood in which they reside [43]. Different segments of the population could indeed be exposed to a heterogeneous risk of infection and disease depending on their age, occupation (e.g., construction workers vs. office workers), adequacy of their housing arrangement (e.g., presence of mosquito window screenings and A.C. units), and the built environment in their neighborhood (e.g. presence of abandoned lots/buildings that may provide more breeding areas). A segment of the population that is at particular risk of arbovirus infection is represented by people experiencing unsheltered homelessness as they are exposed nearly 24/7 to mosquito vector bites. It has been shown that the prevalence of WNV in Houston, Texas, is higher in people that have been homeless for more than one year when compared with the general population [44]. In other metropolitan areas in the United States, including Maricopa County, Arizona, it is so far unknown to what extent vulnerable and underserved populations are being disproportionally exposed to arboviruses. These groups are already affected by large disparities in terms of mortality and a broad array of other health outcomes [45], and arriving at a better understanding of the nature and extent of their disproportionate exposure to arboviral infections is crucial for designing policy interventions that can mitigate such a preventable health disparity and advance health equity.

Effective mosquito control operations rely on multiple factors, with the most important being consistent and reliable mosquito surveillance systems, community engagement, and targeted source reduction. Now, more than ever, effective mosquito control strategies are needed to deal with a flux of importation of travelers carrying arboviruses to prevent local outbreaks, and to curtail them when they start. Effective mosquito surveillance systems are key for guiding and implementing mosquito control strategies and should be considered a priority for local, state, and federal public health professionals.

---

**Supplementary Table 1. Number of traps and mosquitoes collected by the Maricopa County Environmental Services Department Vector Control Division Surveillance System from 2011 to 2021**.

**Supplementary Table 2. Number of mosquitoes (*Aedes aegypti, Culex quinquefasciatus*, and *Culex tarsalis*) collected per trap**.

**Supplementary Table 3. Number of mosquitoes (*Aedes aegypti, Culex quinquefasciatus*, and *Culex tarsalis*) collected by the same set of 478 traps in use since 2016**.

**Supplementary Figure 1. Kernel density-based heat map of the relative abundance of vector mosquito species collected by the Maricopa County Environmental Services Department Vector Control Division Surveillance System from 2012 to 2021**. The color gradient represents the sum of mosquitoes collected by each trap in a 12-month period; (A) *Aedes aegypti*; (B) *Culex quinquefasciatus*; and (C) *Culex tarsalis*. The figure was produced using ArcGIS 10.2 (Esri, Redlands, CA), using layers available at the Maricopa County GIS Mapping Applications available at: https://www.maricopa.gov/3942/GIS-Mapping-Applications.

## Disclaimer

The information provided and views expressed in the publication do not necessarily reflect the official views of the Centers for Disease Control and Prevention (CDC).

