## Supplementary figures and images for "Spatiotemporal Distribution of Vector Mosquito Species and Areas at Risk for Arbovirus Transmission in Maricopa County, Arizona"

### Supplemental Figure 1

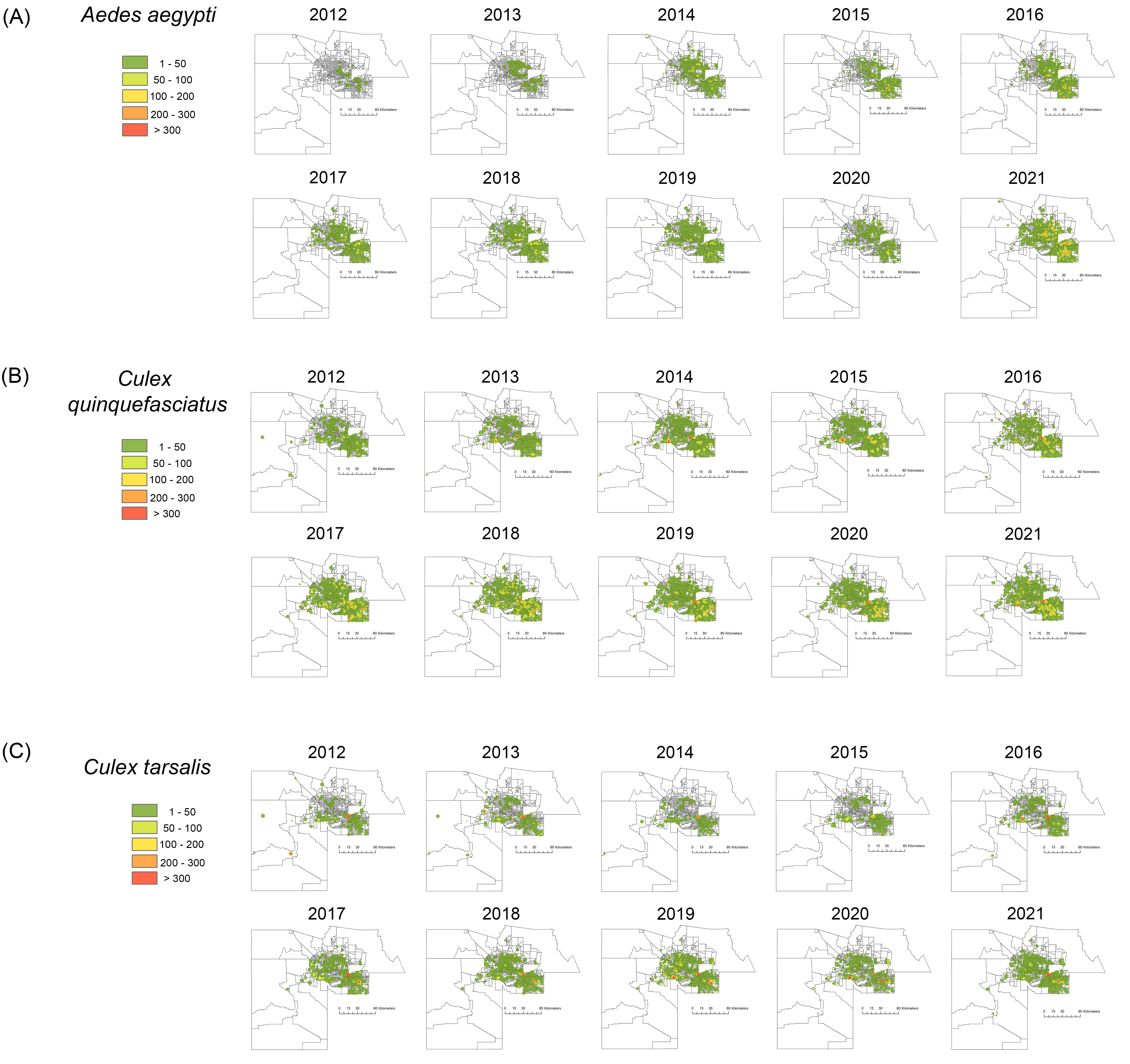
